# Computational investigation for modelling the protein-protein interaction of TasA_(28-261)_ – TapA_(33-253)_: a decisive process in biofilm formation by *Bacillus subtilis*

**DOI:** 10.1101/2020.04.26.062109

**Authors:** Nidhi Verma, Shubham Srivastava, Ruchi Malik, Jay Kant Yadav, Pankaj Goyal, Janmejay Pandey

## Abstract

Biofilms have significant role in microbial persistence, antibiotic resistance and chronic infections; consequently, there is a pressing need for development of novel “anti-biofilm strategies”. One of the fundamental mechanisms involved in biofilm formation is protein-protein interactions of ‘amyloid like proteins’ (ALPs) in extracellular matrix. Such interactions could be potential targets for development of novel anti-biofilm strategies; therefore, assessing the structural features of these interactions could be of great scientific value. Characterization of biomolecular interaction with conventional structure biology tools including X-Ray diffraction and Nuclear Magnetic Resonance is technically challenging, expensive and time-consuming. In contrast, modelling such interactions is time-efficient, economical and might provide deeper understanding of structural basis of interactions. Therefore, during the present study, protein-protein interaction of TasA_(28-261)_–TapA_(33-253)_ (which is a decisive process for biofilm formation by *Bacillus subtilis*) was modeled using *in silico* approaches viz., molecular modelling, protein-protein docking and molecular dynamics simulations. Results identified amino-acid residues present within intrinsically disordered regions of both proteins to be critical for interaction. These results were further supported with PCA and FEL analyses. Results presented here represent novel finding and we hypothesize that aa identified during the present study could be targeted for inhibition of biofilm formation by *B. subtilis*.

## Introduction

Biofilms represent microbial communities embedded within an extracellular matrix consisting primarily of polysaccharides, proteins or extracellular DNA (Costerton, Lewandowski, Caldwell, Korber, & Lappin-Scott, 1995; Kolter & Greenberg, 2006). Several findings have clearly demonstrated that biofilms play a critical role in adaptive microbial processes such as sporulation, resistance against environmental stress etc. (Branda, Vik, Friedman, & Kolter, 2005; Jefferson, 2004). Noticeably, biofilms have been also recognized as a vital element in microbial resistance to antibiotics, microbial evasion of host immune response and establishment of chronic microbial infections(Lynch & Abbanat, 2010). According to a recent report published by National Institutes of Health (NIH), USA, nearly 80% of total chronic microbial infections involve biofilms forming microbial pathogens (Jamal et al., 2018). The process of biofilm formation consists of a rather complex cascade of biomolecular reactions. It initiates with attachment of bacterial cells to a suitable adhesion surface to be followed up with secretion of extracellular matrix giving rise to three-dimensional structures and ending up with a regulated disintegration (Jamal et al., 2018).

Biofilm associated extracellular matrix is largely composed of polysaccharides, extracellular genomic DNA and proteins (Branda et al., 2005; H.-C. Flemming & Wingender, 2010). The exo-polysaccharide components of biofilms are either synthesized extracellularly or intracellularly and subsequently secreted into the outside microenvironment (H. C. Flemming, Neu, & Wozniak, 2007). These polysaccharides serve as the scaffolds for other ECM components viz., proteins, nucleic acids and lipids to adhere (H. C. Flemming et al., 2007). The ECM associated proteins represent a variety of structures and functions including type IV pili, flagella and amyloid-like proteins (ALPs) such ‘curli protein’ in *Escherichia coli* (Chapman et al., 2002), chaplains in *Streptomyces coelicolor* (Claessen et al., 2003), FapC in *Pseudomonas aeruginosa*(Dueholm et al., 2010) and TasA in *Bacillus subtilis (Romero, Aguilar, Losick, & Kolter, 2010*). Characteristically, amyloids like proteins (ALPs) have been recognized as one of the most important components determining the assembly and integrity of biofilm matrix (Romero et al., 2010; Taglialegna, Lasa, & Valle, 2016), yet the corresponding molecular and more specifically the structural information for many of these proteins is rather incomprehensible. Additionally, information pertaining to their interaction with other auxiliary proteins, which are critical for the formation of the biofilm assemblage is also quite inadequate. For example, interaction of processed, extracellular forms of TasA – TapA has been reported to be critical for biofilm formation in *B. subtilis (Romero, Vlamakis, Losick, & Kolter, 2011, 2014*), however, the structural characteristics of this interaction is only poorly understood.

In recent years, *B. subtilis*, a gram-positive, motile, soil-dwelling bacterium has emerged as the model organism of choice for studies pertaining to bacterial biofilm formation (Lemon, Earl, Vlamakis, Aguilar, & Kolter, 2008; Vlamakis, Chai, Beauregard, Losick, & Kolter, 2013). According to the available literature, the processed form of TasA, which lacks 27 amino acid residues at the N-terminus, (Referred as TasA_(28-261)_ hereafter), is the main proteinaceous component of *B. subtilis* biofilm (Branda, Chu, Kearns, Losick, & Kolter, 2006). It has been shown to assemble in amyloid-like fibrillary structure that is critical for holding the biofilm together (Romero et al., 2010). A functional TasA operon has been reported to be critical for biofilm formation in *B. subtilis;* it has been previously reported that a frame mutation in one of the three genes of TasA encoding operon results in a defective biofilm matrix formation (Branda et al., 2006). Similarly, other studies have shown that TasA defective mutant is not able to form biofilm but produce cell bundles that were not attached to each other (Mielich-Süss & Lopez, 2015).

During the biofilm formation in *B. subtilis*, the soluble monomeric form of TasA undergoes proteolytic cleavage leading to conversion of the soluble form TasA into protease-resistant, ß sheet rich amyloid-like protein TasA_(28-261)_. Noticeably, this conversion decisively depends upon the activity of two auxiliary proteins viz., SipW and TapA (Romero et al., 2011; Terra, Stanley-Wall, Cao, & Lazazzera, 2012). SipW, a type 1 signal peptidase, is essential for processing of globular TasA (i.e. removal of 27 amino acid sequences - signal peptide from the N-terminus and release of mature TasA_(28-261)_) and its transport to the extracellular matrix (Serrano et al., 1999; Terra et al., 2012). SipW also catalyzes the proteolytic cleavage and processing of TapA resulting in the release of 30 amino acid signal sequence of mature TapA_(33-253)_ (Terra et al., 2012). TapA_(33-253)_ is the other protein critical for anchoring the fibers to the bacterial cell wall and assemblage of mature TasA_234_ into amyloid-like fibers mature TapA_223_ is (Romero et al., 2011). Collectively, the whole cascade i.e. (i) the processing of globular TasA and TapA; and (ii) mature TapA_223_ mediated anchoring, an assemblage of mature TasA_234_ in amyloid-like fibers are critical for the formation of *Bacillus* biofilm. Furthermore, it has also been reported that the mature TapA_223_ constitutes a minor component of the TasA_234_ fibers in the extracellular matrix(Romero et al., 2014).

In light of the significance of the TasA, SipW, and TapA in biofilm formation by *B. subtilis*, the structures of each of these proteins and their molecular interaction must be vital information and resource towards the development of Anti-*Bacillus* Biofilm Strategy. However, until very recently, the information about the structural characteristics of these molecules and their interactions was relatively obscure. The crystal structure of the soluble monomeric form of TasA was reported only a few months back(Diehl et al., 2018). It revealed that the monomeric soluble unprocessed form of TasA mainly consists of a jellyroll fold composed of 2 antiparallel ß-sheets flanked by 6 short helices and a long loop region(Diehl et al., 2018). During processing of TasA, its structure undergoes a transition of its folded form to protease-resistant biofilm-stabilizing fibrils(Diehl et al., 2018).

Several observations from past reports indicate that the interactions of the processed forms of TasA and TapA (referred as TasA_234_–TapA_223_ hereafter) are potentially a critical factor in the formation of functional *Bacillus* biofilm(Romero et al., 2014). Therefore, we hypothesize that the interference of this interaction could be exploited for the development of an *‘anti-Bacillus* biofilm strategy’. With this hypothesis, we carried out Computational analyses of *B. subtilis* TasA_234_ - TapA_223_ interaction using homology modeling, rigid protein-protein docking, and molecular dynamics approach. In particular, we analyzed the protein-protein docking confirmations for revealing the critical amino–acid residues involved in the interactions.

## Methods and Materials

### Sequence retrieval and molecular characterization

The protein sequences of TasA and TapA of *B. subtilis* subspecies subtilis strain 168 were obtained from National Centre for Biotechnology Information (NCBI) protein database www.ncbi.nlm.nih.gov/ with accession numbers NP_390342 and CAB14395 respectively(Coordinators, 2013).

### Homology modeling of TasA_234_ and TapA_223_ and model validation

The crystal structure of proteins being analyzed within this study was not available, therefore, the amino acid sequence of target proteins (i.e. TasA_234_ and TapA_223)_ were subjected to three dimension structure prediction using homology modeling strategy. For a selection of homology modeling templates, PDB-BLAST analyses were carried out using the primary amino acid sequences of the target proteins as the query sequence. From this analyses, proteins showing highest sequence identity and query coverage were selected as a template for model building using Modeller version 9.14 (https://salilab.org/modeller/9.14/release.html)(Eswar et al., 2006). The sequence alignments were derived and used for 3-dimensional structure predictions by the implication of restraints from the template structures and their alignment with targeted protein sequences. The predicted 3-D models were further refined by DOPE (discrete optimized protein energy) score assessment method in Modeller version 9.14. The quality of the modeled structure with respect to stereo-chemical geometry and energy were analyzed by PSVS (protein structure and validation suite) including Procheck (Laskowski, MacArthur, Moss, & Thornton, 1993). The PDB structures were also used for further validation of the predicted models of mature TasA and TapA as well as during *in silico* docking and dynamic studies (described later).

### Prediction of disorder region and sequence-based interaction analysis

According to the available literature, the mature forms of TasA and TapA (i.e.TasA_234_ and TapA_223)_ have the characteristic properties of Amyloid-Like Proteins; therefore, they may have the presence of intrinsically disordered regions. With this rationale, *in silico* prediction of the disorder regions was carried out with TasA_234_ and TapA_223_. For this, an online tool (i.e. PONDR; Predictor of Natural Disordered Regions available at http://www.pondr.com/) (Xue, Dunbrack, Williams, Dunker, & Uversky, 2010) was used. It uses the combination of six predictors (VLXT, VL3-BA, VSL2, XL1_XT, CAN_XT, and CDF) for the prediction of each and every residue along the sequences. It predicts disordered and ordered residues by maximizing the number of training-set disordered residues while simultaneously maximizing the number of training-set ordered residues giving outputs 0.5.

The primary amino acid sequence based interaction study of TasA_234_ and TapA_223_ was carried out by using STRING 10 database (https://string-db.org/) (Szklarczyk et al., 2017) which includes both direct (physical) and indirect (function) interactions derived from scientific literature, co-expression analysis, and gene ontology.

### Molecular dynamics simulations of modeled TasA_234_ and TapA_223_

The molecular dynamics simulation of TasA_234_ and TapA_223_ models and their interactions during formation of different dock complexes were performed by using Desmond (Desmod molecular dynamics system, D.E. Shaw Research, NewYork, NY, 2019) and running on a Bio-Linux cluster (64-bit processor). For these analyses, the predicted 3D structures were stabilized by using default parameters with minor variations. An ‘all-atoms optimized potential liquid for simulations (OPLS-AA) force field was use and all the simulations were conducted in the SPC water environment, i.e. a ‘bt’ cubic box, SPC water model and neutralized by the addition of positive Na^+^ and negative Cl^-^ ions. The energy minimization was performed for 5,000 cycles by using the steepest descent method. Furthermore, temperature and pressure equilibration were performed by heated to 300K (= ~ 27°C) for 100 ps i.e, NVT using Berendsen thermostat algorithm and 100 ps steps NPT by using Nose-Hoover thermostat respectively. The final production run was performed for 100ns for each system using the same pressure, temperature and integrator. Molecular dynamics trajectories were then visualized and analyzed using the Visual Molecular Dynamics (VMD) (https://www.ks.uiuc.edu/Research/vmd/) (Humphrey, Dalke, & Schulten, 1996). All the graphs of different simulations were plotted using GraphPad Prism version 7.0 (https://www.graphpad.com/)(Prism, 1994).

### Protein-protein docking & molecular dynamics studies for prediction interaction

To predict the physical interaction(s) between TasA_234_ – TapA_223_, the protein-protein docking studies was performed using rigid docking prediction server viz., Z-DOCK (http://zdock.umassmed.edu/) (Pierce et al., 2014). It predicts interaction(s) of protein-ligand and protein-protein on the basis of an FFT (Fast Fourier Transform) algorithm. Additionally, it also explores a variety of docked lower energy conformations using shape complementarily and electrostatic potential. The protein-protein interactions were also evaluated with another online protein-protein interaction prediction server viz., ClusPro (https://cluspro.org/help.php) (Kozakov et al., 2017). It ranks the interactions according to the best cluster size and four different sets of energy coefficients. This server generates putative interaction complexes taking into consideration some of the critical features of protein-protein interaction e.g. electrostatic, hydrophobic and Van der Wall interactions. The resulting best interaction - complexes were ranked according to the lowest free energy and complexes were refined over 10 ns time scale using the above mentioned operational parameter. The docked complexes from ClusPro were then further used by another protein-protein docking server FireDock (bioinfo3d.cs.tau.ac.il/FireDock/) (Mashiach, Schneidman-Duhovny, Andrusier, Nussinov, & Wolfson, 2008) to optimize and re-score the complexes by restricting protein flexibility of the interacting surfaces.

For determining the dynamic stability of the protein-protein interaction complexes(s) generated with different servers, the molecular dynamics studies of dimeric complexes(s) were performed using Gromacs 2016.5. The protein-protein interactions of docked complexes were examined by using InterProSurf web server (http://curie.utmb.edu/prosurf.html)(Negi, Schein, Oezguen, Power, & Braun, 2007) and PyMol (https://pymol.org/2/)(DeLano, 2002).

### Contact map analysis

The putative binding site(s) and interfaces in homo-molecular and hetero-molecular complexes were confirmed by analyses of intermolecular contact maps determined by COCOMAPS (https://www.molnac.unisa.it/BioTools/cocomaps/)(Vangone, Spinelli, Scarano, Cavallo, & Oliva, 2011). It provides interactive contact maps from the 3D biomolecular complexes on the basis of distance range and physicochemical properties e.g. hydrophobic, hydrophilic interactions. During our study, the stabilized interaction complexes with the lowest energy and the highest conformation stability obtained after molecular dynamics were used for contact map analyses.

### Principal component analysis (PCA) and Free energy landscape (FEL)

The principal component analysis (PCA) was carried out to systematically represent the linear transformation of characteristics of the covariance matrix and reduction of the instantaneous linear correlations among the coordinates of the biomolecules. This analysis was performed with GROMACAS –version 2019(Pronk et al., 2013) according to the method described earlier(Amadei, Linssen, & Berendsen, 1993; Ichiye & Karplus, 1991). During the present study, fluctuations were assessed by selecting Cα atoms of target proteins, which provides significant characteristics to essential internal motions. Results obtained from PCA analyses were subjected to free energy landscape assessment. The g_anaeig and g_covar utilities were used for revealing the changes in the motion patterns of the protein complexes. The g_sham utility for FEL (free energy landscape) analysis was used to estimate the joint probability distribution on to the three-dimensional space defined by first two principal components, i.e., PC1 and PC2.

## Results

### Homology modeling of TasA_234_ and TapA_223_ and model validation

The primary amino acid sequences of both the proteins (TasA & TapA; retrieved from NCBI) were subjected to PDB BLAST analyses. Results from the PDB BLAST analyses showed that TasA has the highest sequence similarity with a conserved exported protein from *Bacteroides fragilis* (PDB ID: 3CLW)(Halgren, 2009) and D-isomer specific 2-hydroxy acid dehydrogenase from *Lactobacillus delbrueckii* ssp. Bulgaricus (PDB IDs: 2YQ4)(Holton, Anandhakrishnan, Geerlof, & Wilmanns, 2013). TapA showed highest sequence similarity with a Conserved Protein of Unknown Function from *Staphylococcus aureus* (PDB IDs: 2AP3)(Costerton et al., 1995). The amino acid sequences of best-matched PDB structures were selected as template for further studies and retrieved from NCBI and subjected to the off-line Alignments with the sequences of TasA and TapA using CLUSTAL Omega(Sievers et al., 2011). The alignment reports obtained from CLUSTAL Omega analyses are presented in Supplementary figure S1A, S1B. Based on these observations, the respective PDB structures were selected as template structures for homology modeling of TasA, TapA and their respective processed mature forms (i.e. TasA_234_ and TapA_223)_. Twenty 3-dimensional models each for TasA_234_ and TapA_223_ were generated with homology modeling. Each of them was found to have a characteristic unique “Discreet Optimized Protein Energy (DOPE) scores. The models were compared for their DOPE scores and best models were selected on the basis of lowest DOPE scores. Herein, models with DOPE scores of −56877.059 and −18539.964 were found to be the best models for TasA_234_ and TapA_223_ respectively. These models were selected and subjected to quality assessment, model refinements. The refined models were validated with Ramachandran Plot Analyses. Results from this analysis showed that, for the selected models, most of the residues lie in the favored, allowed, and generously allowed regions. Only 3.5% and 0.0% residues were found to lie within in the disallowed region in case of models of TasA_234_ and TapA_223_ respectively.

### Stability Analyses of protein models with Molecular Dynamics Simulations

The models of TasA_234_ and TapA_223_ selected and validated above were analyzed for their stability by performing molecular dynamics simulations performed over a time scale of 100 ns. During MD simulation of TasA_234_, nearly all the residues remained in the most favored or additionally allowed or generously allowed regions and only 0.9% of residues were found to be in the disallowed region. The structural integrity of most of the regions was sustained during molecular dynamics except for the residues Glu-50, Ser-77, Val-116, Leu-140, Tyr-181, His-242, and Asn-252 (Fig 1A, B). Similarly, in case of TapA_223_, only 1.3% residues were found to remain in the disallowed regions as indicated by the structural integrity most of the regions except Asp-40, Asp-60, Cys-92, Glu-116, and Glu-231 (Fig 1C, D).

**Figure 1:**
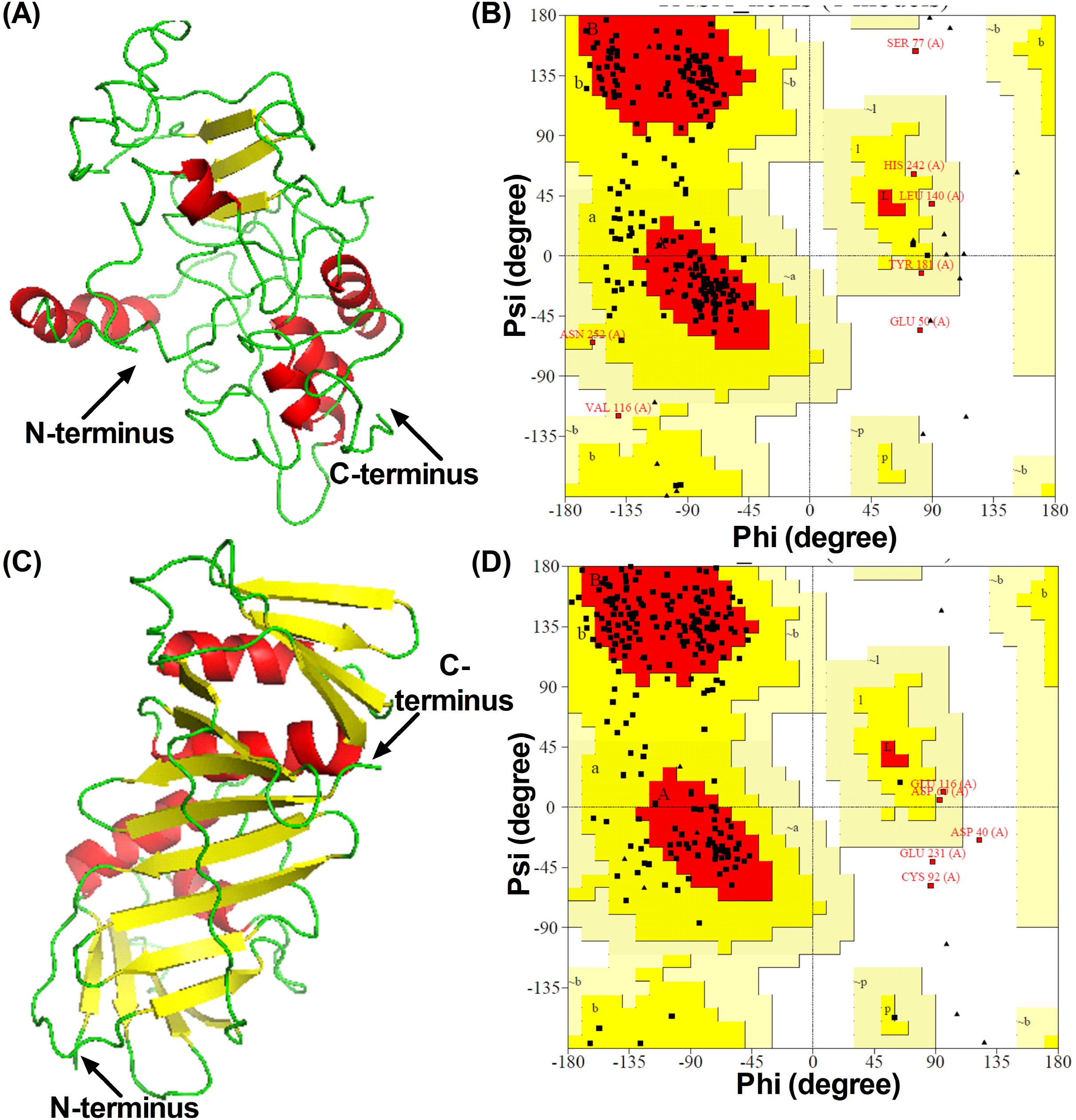
Protein 3-D modeling and Validation: (A, C) the 3-D model of TasA and TapA build by Modeller 9.14. The refined and stabilized 3-D structure of proteins after MD at 100 ns have visualized in Pymol. The cartoon structure represents the secondary structural elements are shown helices (red), ß-sheets (yellow) and coils (green)in color. (B, D) Ramachandran plot analysis by using Procheck, indicating residues in the favored regions, allowed regions, generously allowed regions and disallowed regions have shown by red, yellow, light yellow and white color respectively.

To further assess the stability of generated models for TasA234 and TapA223, results obtained from molecular dynamics simulations were also subjected to position restraint for 100 ns. The Root Mean Square Deviation (RMSD) of backbone atoms were calculated over the entire simulation trajectories. The RMSD Plots of modeled proteins showed that the stability of structures was maintained even after 100 ns of MD simulation. The RMSD values were found to range between 0.5 to 1 nm for TasA234 and 0.5 to 0.8 nm for TapA223 (Fig 2A, 2B). Similarly, the Root Mean Square Fluctuation (RMSF) of Cα atom for modeled proteins was also evaluated and the flexibility of each residue was determined during the simulation time of 100 ns. The overall fluctuation rates of TasA234 and TapA223 model structures remained in the range of 0.5 to 1nm and 0.5 to 0.8 nm respectively (Fig 2C, 2D). Thus, it could be inferred that during MD simulations, the modeled structures of both TasA234 and TapA223 remain in stable configurations. One of the noteworthy observations during MD simulations was that the residues present at C-terminal of TasA234 were found to be more flexible compared to other regions of the modeled protein (Fig. 2C). In the case of TapA223, the region close to C-terminal domain is quite flexible in comparison to the N-terminal domain. The region close to N-terminal domain exhibits significantly lower fluctuations (Fig. 2D).

**Figure 2:**
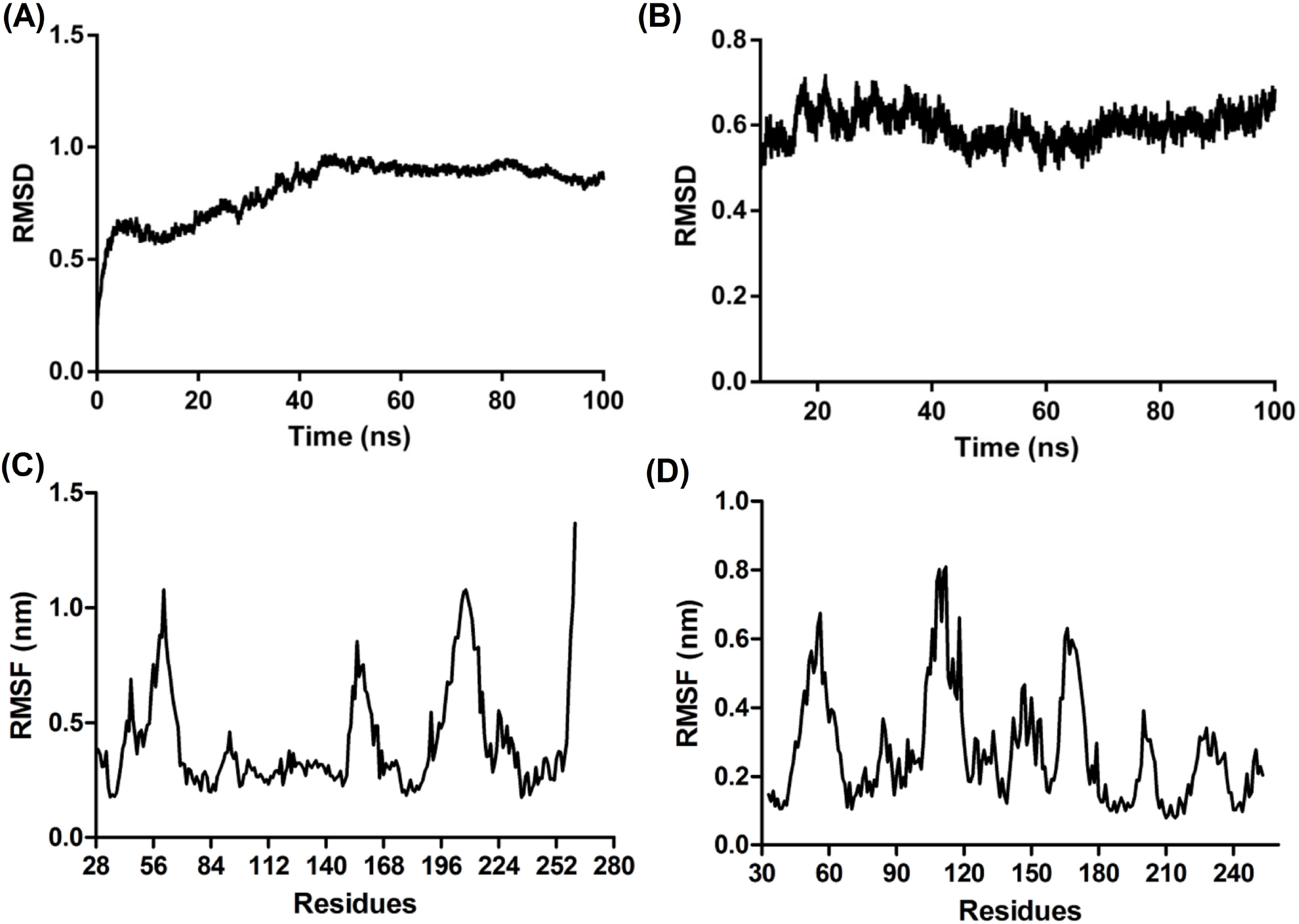
Molecular dynamic study of TasA and TapA: (A, B) RMSD calculations of TasA and TapA over the entire MD simulation trajectories at 100 ns. (C, D) RMSF evolution of TasA and TapA represents the flexibility of each residue during the simulation time.

This observation is consistent with the recent report which described TapA to be composed of two interacting domains; a disordered C-terminal domain and a more structured N-terminal domain(Abbasi et al., 2018). Noticeably, the two-domain composition of TapA has been suggested to be important for its interactions in the extracellular matrix in biofilms(Abbasi et al., 2018).

### Assessment of the disordered regions in TasA_234_ and TapA_223_

With the rationale that the intrinsically disordered regions of proteins contribute to their interactions with other proteins, an analysis was carried out to predict the disordered region in target proteins. Results from this analysis showed that there are four stretches (i.e. amino acid residues 42-71, 116-126, 150-200 and 241-261) that could potentially constitute the intrinsically disordered regions in case of TasA_234_. In the case of TapA_223_, the disordered region spreads in form of a single stretch from amino acid residues 200-251 (Supplementary Fig S2A & S2B). Again, these results were consistent with the report describing the disordered domain close to C-terminus in case of TapA(Abbasi et al., 2018).

### Protein-protein interaction analyses

In order to predict the protein-protein interactions of the target proteins, the ‘String Analysis’ was performed using primary amino acid sequences. Results from this analysis showed that TasA might interact with 10 different proteins (Supplementary Fig. S3). Most noticeably, amongst the predicted interacting proteins, TapA (yqxM) was found to have the highest confidence score of 0.996 (Supplementary Fig S3). This analysis clearly recognized that TasA and TapA have strong protein-protein interactions with one other. This observation is in agreement with the previous report, where processed TapA has been reported as an essential accessory protein interacting with processed TasA and contributing to the polymerization of TasA_234_ into an amyloid fiber and biofilm formation(Romero et al., 2011, 2014).

### Protein-Protein docking of TasA_234_-TapA_223_

To explore the specific TasA_234_-TapA_223_ interactions the molecular docking analyses for protein-protein interactions were performed using a rigid body docking program (viz., Z-DOCK, CLUSPRO). These analyses resulted in the generation of several thousands of the complexes. Top 10 complexes for each interaction were selected and later subjected to MD simulations at 10 ns for minimization and optimization of their geometry. The results obtained from rigid docking analyses were later validated by use of a flexible refinement program (i.e. FireDock).

### Molecular Interactions characterized by Z-DOCK

The interaction of TasA_234_-TapA_223_ in the docked complex showed the formation of hydrogen bonds between ND2-atom of Asn54 and SG atom of Cys92; NZ atom of Lys65 and O atom of Val91; OD1 atom of Asn54 and NZ atom of Lys88; OG1 atom of Thr177 and N atom of Ser82. The detail of hydrogen bond interactions of TasA_234_-TapA_223_ are listed in table 1

**Table No.1:** The protein interface residues of TasA-TapA complex formed by Z-Dock server participate in hydrogen bond formation.

| ACCEPTOR |  |  |  | DONOR |  |  |  | Dist (Å)<br>H-Bonds |
| --- | --- | --- | --- | --- | --- | --- | --- | --- |
| Res<br>Code1 | N°<br>res1 | Atom1 | Chain1 | Res<br>Code2 | N° res2 | Atom2 | Chain2 |  |
| ASN | 54 | ND2 | A | CYS | 92 | SG | B | 1.88 |
| LYS | 65 | NZ | A | VAL | 91 | O | B | 2.98 |
| LYS | 88 | NZ | B | ASN | 54 | OD1 | A | 2.71 |
| SER | 82 | N | B | THR | 177 | OG1 | A | 2.36 |

### Molecular Interactions characterized with ClusPro 2.0

Molecular interactions within docked complexes were also analyzed with another rigid docking program viz., ClusPro 2.0. The complexes were recollected and analyzed on the basis of best electrostatic and desolvation free energies. The details of interaction characteristics are listed in Table 2.

**Table No.2:** Docking energy values for hetero-molecular TasA234 and TapA223 using ClusPro.

| Complex | TasA239-TapA223 |  |
| --- | --- | --- |
|  | Center | lowest energy<br>(in kcal/mol) |
| 0 | -635.9 | -878.6 |
| 1 | -596.5 | -763.5 |
| 2 | -740.6 | -741.9 |
| 3 | -635.5 | -808.1 |
| 4 | -603.8 | -703.6 |
| 5 | -593.8 | -860.7 |
| 6 | -586.5 | -787.3 |
| 7 | -612.5 | -694.6 |
| 8 | -617.6 | -720 |
| 9 | -655.6 | -707.3 |

The TasA_234_-TapA_223_; best-docked complex showed a cluster size 155 having lowest energy - 878.6. The hydrogen bonds formation was observed between NE2-atom of Gln51 and O atom of Ala96; ND2 atom of Asn56 and O atom of Lys88; OG2 atom of Ser58 and OE1 atom of Glu56; NZ atom of Lys65 and O atom of Thr90; NZ atom of Lys144 and OD2 atom of Asp83; NZ atom of Lys147 and OE2 atom of Glu116; OD1 atom of Asp31 and N atom of Thr90; OE1 atom of Gln51 and NE2 atom of Gln62; O atom of Asn54 and SG atom of Cys58; O atom of Leu57 and SG atom of Cys58; O atom of Gly183 and N of Lys119; OD1 atom of Asp241 and NE atom of Arg124.

### Molecular Interactions characterized by Fire-DOCK

In order to further examine the protein-protein interactions predicted by rigid docking, the flexible refinement analyses were performed for various complexes of TasA_234_ and TapA_223_ using Fire-DOCK program. The complexes were selected on the basis of the lowest global energy values (Kcal/mol). From this analysis, the best TasA_234_-TapA_223_ docked complex exhibited maximum global energy value of - 90.19 Kcal/mol. The detail of global energies of docked complexes is presented in table 3. A list of all the residues predicted for involvement in the interaction during formation of TasA_234_-TapA_223_ complexes is presented in table 3. The noticeable observation with these analyses was that none of the hydrogen bonds were common with respect to protein-protein interaction.

**Table No.3:** Fire Dock docking energy values for TasA234 and TapA223 complex; where ACE and HB stand for atomic contact energy and hydrogen bonds respectively.

| Fire Dock energies | TasA<sub>234</sub>-TapA<sub>223</sub> |
| --- | --- |
| Global energy | -90.19 |
| Attractive Vdw | -63.39 |
| Replusive Vdw | 10.41 |
| ACE | -12.07 |
| HB | -4.1 |

### Molecular Interactions analyses after MD simulations

The complexes analyzed with rigid and flexible docking were also subjected to energy minimization and optimization by MD simulation studies at the time scale of 100 ns. The structural integrity of the complexes in terms of RMSD and RMSF showed that the complexes were stable during the entire MD simulation trajectories for TasA_234_-TapA_223_ complexes. The Interaction analysis of TasA_234_-TapA_223_ reveals that H-bonding interactions play an important role in the formation and stabilization of dimmers. Noticeably, many of the hydrogen bonds that were previously observed with Z-DOCK and ClusPro analyses remained stable after MD simulation. In addition, a few altered interactions were also observed after completion of the MD simulation. The details of the interacting residues involved in H-bonding, hydrophobic and hydrophilic interactions identified by different docking programs are summarized in table 4.

**Table No.4.** Protein interface residues of TasA_28-261_ and TapA_33-253_ complex participate in hydrogen bond formation (before molecular dynamics simulation studies).The amino acid residues highlighted with ‘red’ color fonts are those identified by all three protein – protein interaction Softwares used during the present study.

| Complexes | Molecules | Interacting residues in Z-DOCK | Interacting residues in ClusPro | Interacting residues in Fire-Dock |
| --- | --- | --- | --- | --- |
| TasA<sub>28-261</sub>-<br>TapA<sub>33-253</sub> | TasA<sub>28-261</sub> | Asn54, **Lys65**,<br>Thr177 | Gln51, Asn56, Ser58,<br>**Lys65, Lys144, Lys147** | Asp31, Gln51, Asn54,<br>Leu57, Ser58, **Lys65**,<br>**Lys144, Lys147**, Asp241 |
|  | TapA<sub>33-253</sub> | Ser82, **Lys88**,<br>Val91, Cys92 | **Cys58, Gln56, Gln62,**<br>**Asp83, Lys88, Thr90,**<br>Ala96, Lys119, Arg124 | **Gln56, Cys58, Gln62, Asp83,**<br>**Thr90**, Glu116, Arg124 |

**Table No.5.** The protein interface residues of TasA(28-261) and TapA(33-253) complexes participate in hydrogen bond formation (After molecular dynamics simulations).

| Complexes | Molecules | Interacting residues in Z-DOCK | Interacting residues in ClusPro | Interacting residues in Fire-Dock |
| --- | --- | --- | --- | --- |
| TasA(28-261)<br>TapA(33- 253) | TasA(28-261) | Ala40, Asn74, Glu103,<br>Lys118, Glu119 | Asp31, **Asn56, Ser58,**<br>**Asn59,** Leu66, **Lys144,**<br>**Lys147,** Arg195 | Asp31, Asn56, Leu57,<br>Ser58, **Lys65, Lys144,**<br>**Lys147,** Arg195 |
|  | TapA(33- 253) | Ser93, Trp113, **Glu116,**<br>Lys119, | **Glu56,** Asp86, **Lys88,**<br>Val91, **Cys92, Glu116** | **Gln56, Cys58, Gln62,**<br>**Asp86, Cys92, Glu116,**<br>Arg124 |

The stereo view of protein interface and interacting residues involved in H-bonding of TasA_234_-TapA_223_ hetero-molecular complexes obtained from molecular simulation analysis of Z-Dock, ClusPro and FireDock shown in fig 3A-C. The RMSF value of Cα atom of each complex was plotted to evaluate the flexibility of each residue during simulation time. The overall fluctuation rate of complex structure was observed to be within the range of 0 to 7Å respectively. The RMSD value of the interaction complex was also found to be in the stable range.

**Figure 3:**
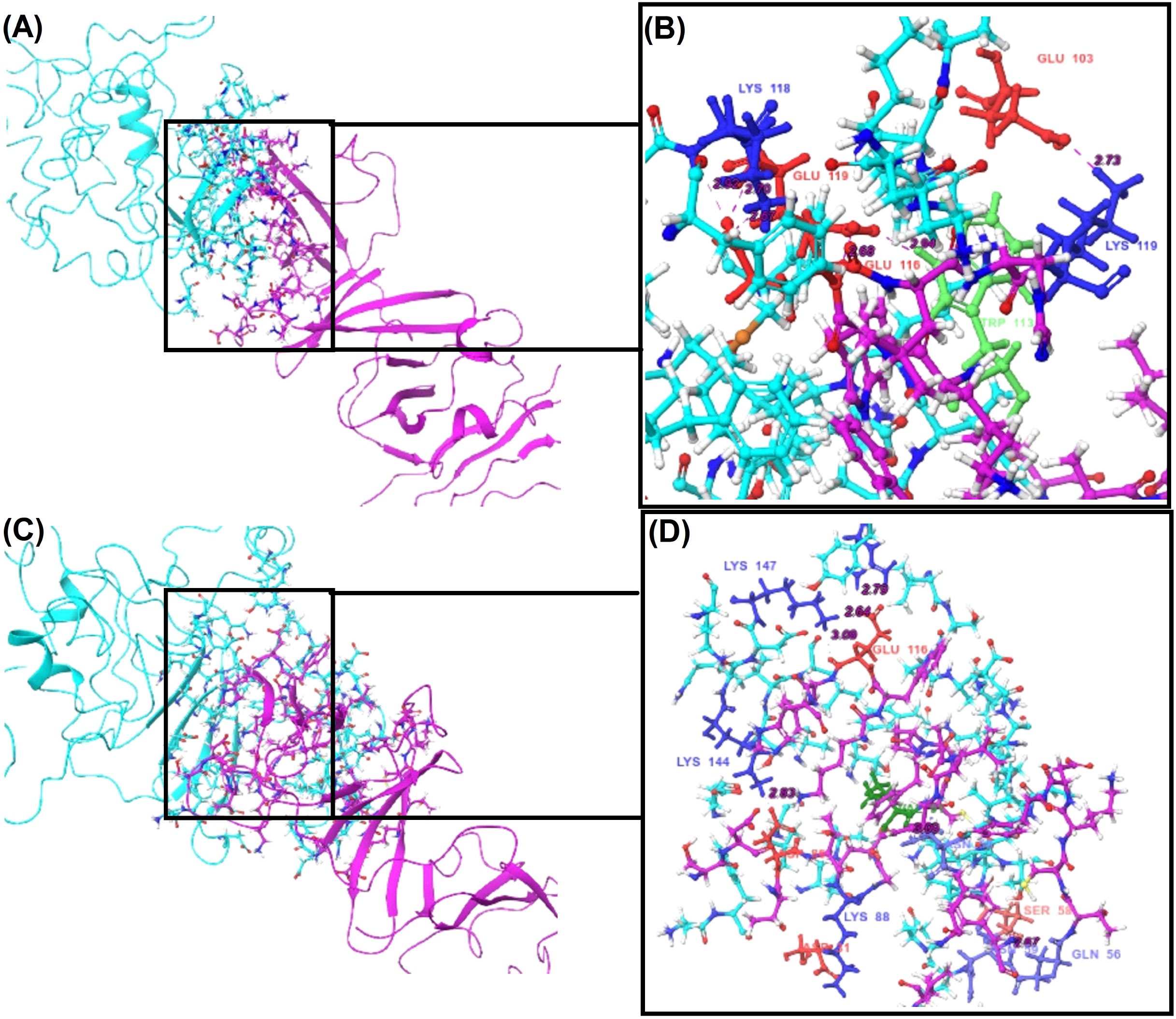
TasA_(28-261)_ - TapA_(33-253)_ interaction analysis: (A, C) The cartoon view of the interface residues of TasA_234_-TapA_223_ complex involve in h-bonding, where ChainA and ChainB represent by green and cyan color respectively visualized in Maestro 11. (B, D) The atoms of interacting residues represent by residue type information and h-bond network between interacting residues represents by dashed lines (pink in color).

The MD simulation analyses of the TasA_234_-TapA_223_ complex obtained with Z-Dock, ClusPro, and Fire-Dock showed that the complexes are stable. The RMSD values of these complexes were found to be 0.000 – 7.63; 0.000 – 9.87; and 0.000 – 8.7 respectively over 100 ns (Fig. 4A). During MD simulations, these complexes showed very little fluctuations as indicated by RMSF values of 1.059 – 7.049; 0.826 – 9.461 and 0.83 – 9.518 nm respectively (Fig. 4B). An increasing trend of fluctuations was also observed during the MD simulation, yet, the RMSD value remained the same and the complexes remained stable Fig 4A-B.

**Figure 4:**
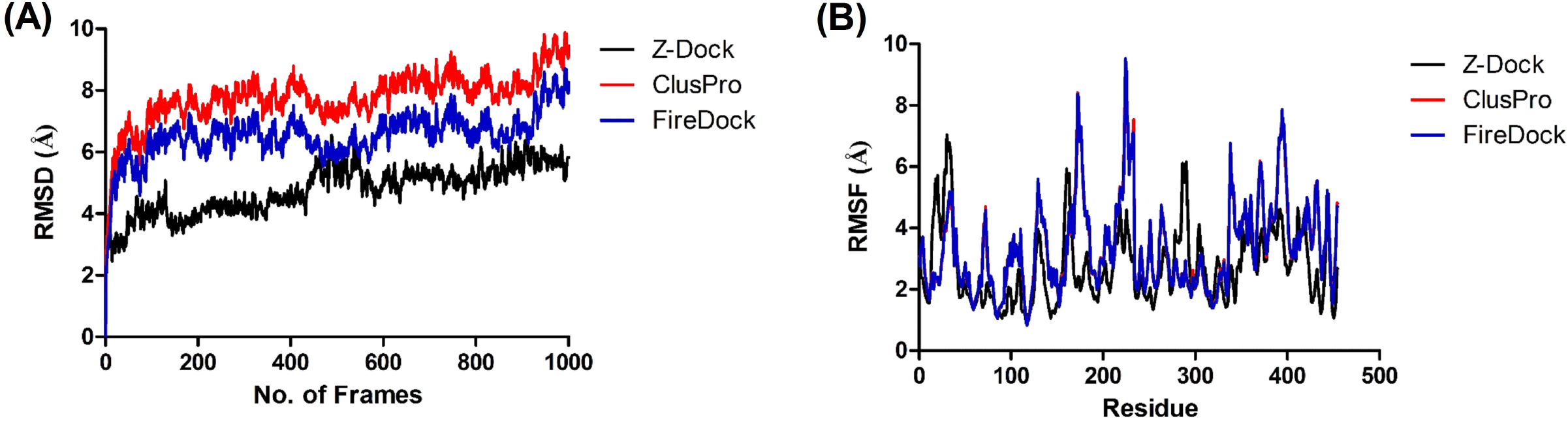
a molecular simulation analysis of TasA_(28-261)_-TapA_(33-253)_ complexes: (A) in this RMSD graph of hetero-molecular docked complexes from different docking programs from the 100ns simulation time frame. B) RMSF calculations of hetero-molecular complexes over the entire MD simulation giving a clear view of fluctuating regions.

### Contact Map prediction for TasA_234_-TapA_223_ interactions

The contact maps for TasA_234_-TapA_223_ hetero-molecular interactions complexes showed contacts at distance ranges and physiochemical properties. The distance ranges in the contact map showed contacts at an increasing distance within 7, 10, 13 and 16 Å (Fig 5). The property contact map showed the physiochemical nature of interacting residues which revealed that 56-74, 103-119, 144-147 and 195 regions of TasA_234_ and 56, 88-92 and116 regions of TapA_223_ are the major interacting residues which participate in interactions of these proteins. This observation is in accordance with the results obtained from protein-protein interaction docking analyses and the subsequent MD simulation analyses. These findings are also in agreement with the previous reports wherein some of the important residues of TasA_234_ and TapA_223_ were biochemically characterized for their potential involvement interaction of these proteins.

**Figure 5:**
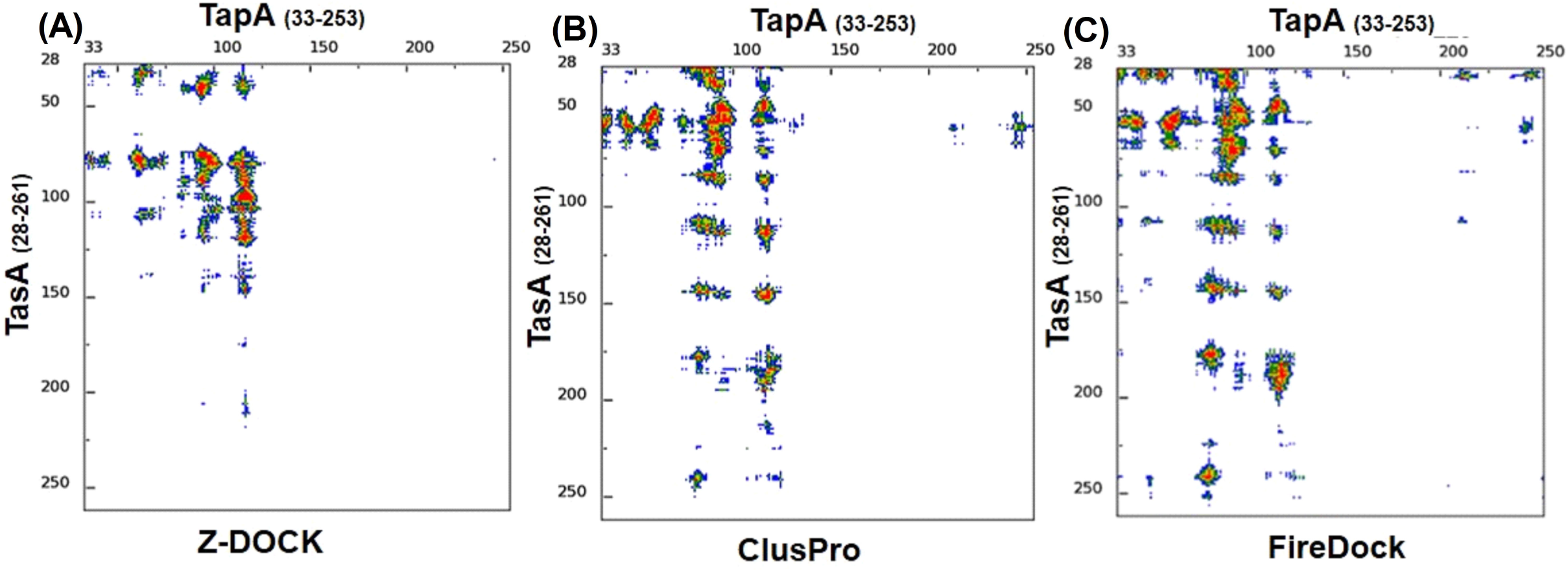
Contact map analysis of TasA_(28-261)_-TapA_(33-253)_ complex at increasing distance range, such that 7Å, 10Å, 13Å and 16Å: Contact map of TasA_(28-261)_ - TapA_(33-253)_ complex obtained from ZDock, ClusPro and FireDock determined by Contact distance range, such that 7Å, 10Å, 13Å and 16Å are represented by orange, yellow, green and blue colors respectively.

### Principal component analysis (PCA) and Free energy landscape Analyses (FEL)

The PCA analysis was performed to correlate the internal motion of different molecules and largest root mean square fluctuations of atoms in biomolecules by using a 3N x 3N matrix of Cartesian displacements. To estimate the conformational transition of TasA_234_-TapA_223_ complexes PCA was performed on data generated from molecular dynamics simulation over 10 ns dynamics trajectories. The atomic motions of TasA_234_-TapA_223_ complexes were analyzed and the coordinates of Cα atoms and characterized by an eigenvector and eigenvalue. The correlated residue motions of TasA_234_-TapA_223_ complexes were plotted by using the covariance matrix (fig 6A). It was observed that the first eigenvalues of all atoms in homo-molecular and hetero-molecular complexes occupy high values; whereas subsequent eigenvalues were observed to be greatly decreased. The atomic motion differences were less observed in docked complex shown in fig (6A-C). Positive regions (red in color) represent the correlated residual movement whereas the negative region (blue in color) indicates the association of anti-correlated residual movement. The anti-correlated motion was dominant in a hetero-molecular complex that might be due to unfolded structures in complexes.

**Figure 6:**
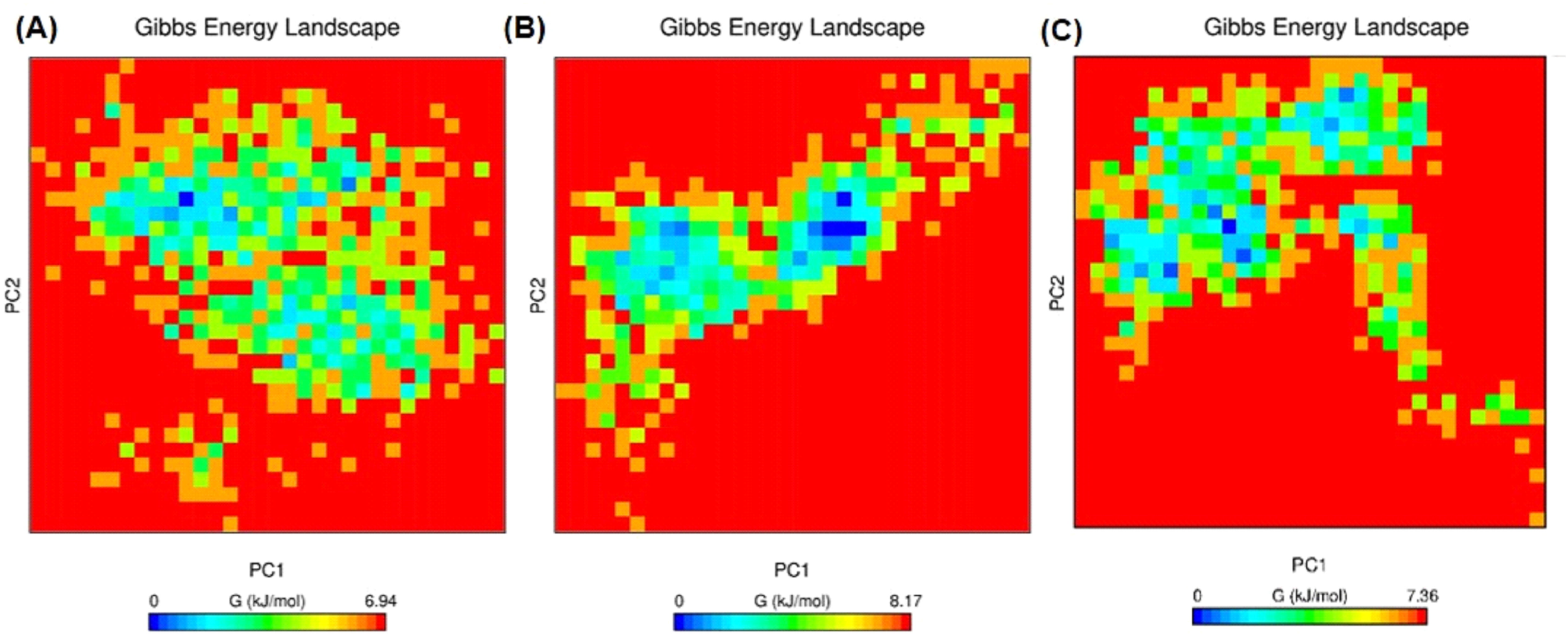
Free Energy Landscape Analysis of TasA_(28-261)_ and TapA_(33-253)_ complex: (A, B and C) show free energy landscape (FEL) between First and Second principal component of TasA_(28-261)_ - TapA_(33-253)_ complex obtained from ZDock, ClusPro and FireDock respectively..

The deviations observed with PCA analyses were further assessed by performing Free Energy Landscape (FEL) analysis. The FEL analysis determines the conformational changes in biomolecules depending upon respective Gibbs free energy acquired during the interaction and complex formation. The blue spots (protein having higher Gibbs free energy) favors an unfolded state and a red spot (decrease in Gibbs free energy) depicts a folded state of biomolecules. From this analysis, it was clearly observed that the TasA_234_-TapA_223_ complex showed higher Gibbs free energy or blue spots. The equilibrated trajectories demonstrated that the TasA_234_-TapA_223_ complexes had significantly high coverage of the eigenvectors as indicated by > 98% of total fluctuations were found to be governed by the initial 10 PCs (Fig. 7A) in comparison to unbound form of TasA_234_ and TapA_223_. The collective motion of bounded and unbounded state of TasA_234_ and TapA_223_ was also provide a clue about the structural changes in proteins (Fig 7B). This observation could be argued to indicate that under natural physiological condition, the TasA_234_-TapA_223_ complexes may be formed more frequently and they attain greater degrees of stability.

**Figure 7:**
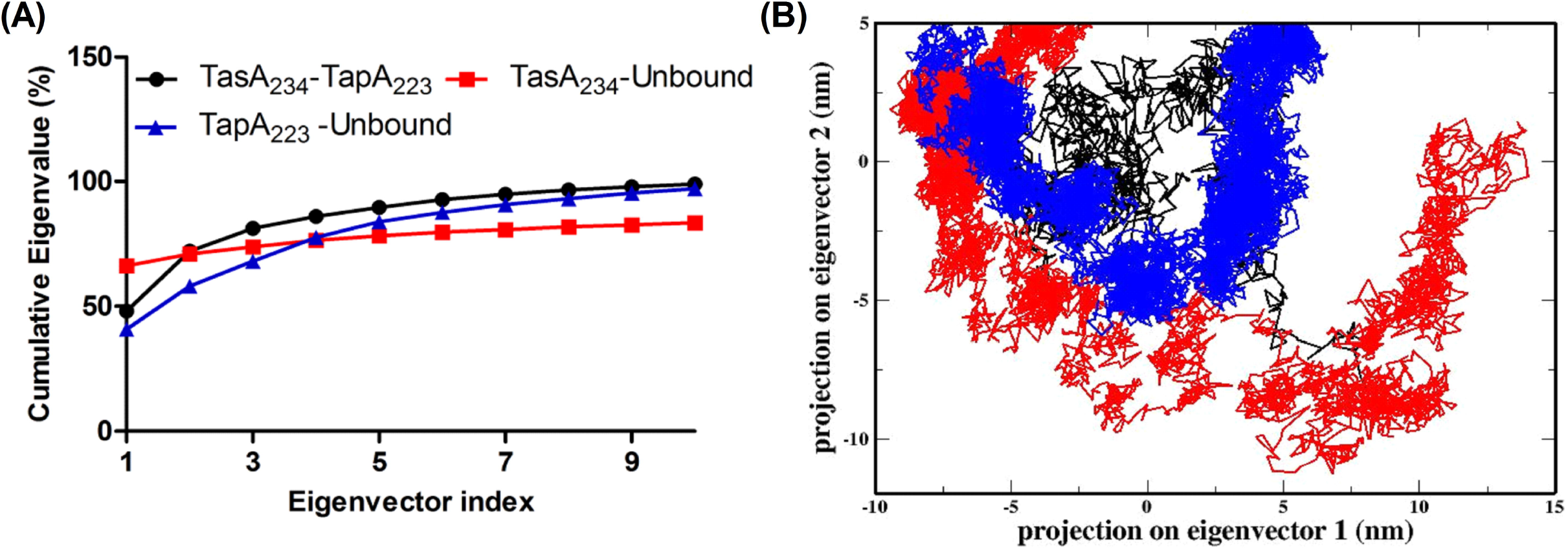
Comparison of bounded and unbounded form of TasA_(28-261)_ and TapA_(33-253)_: (A) represent the comparison first 10 eigenvectors during the simulation time of 10 ns of bounded (black), unbound TasA_(28-261)_ (red) and unbound TapA (blue in color). **(B)** The bounded TasA_(28-261)_ TapA_(33-253)_ (black color) and unbounded TasA_(28-261)_ and TapA_223_ (red and blue color) state represent in phase space along with the first two principal components.

## Discussion

Protein-protein interactions contribute as a vital component of protein functionality. However, studies pertaining to *in vitro* characterization of the structural aspects of such interactions are time-consuming and technically challenging. In the same vein, it is widely acknowledged that the three-dimensional structures of protein-protein interaction complexes are generally more difficult to determine experimentally that the structure of an individual protein(Vakser, 2014). Therefore, the implementation of computational analyses for the characterization of protein-protein interactions is rapidly emerging as a valuable tool for addressing the fundamental questions pertaining to the physiological relevance of the interactions(Vakser, 2014). In the recent past, a number of studies have been undertaken with this approach to characterize the protein-protein interactions using computation approaches. One such study examined the interactions of *Mycobacterium tuberculosis* DprE1 and DprE2(Bhutani, Loharch, Gupta, Madathil, & Parkesh, 2015). Microbial biofilms are important examples wherein protein – protein interactions have pivotal role in defining their integrity and the functionality. Very often, biofilms contain Amyloid-like Proteins (ALPs) which interact with accessory protein(s) during the course of their assembly, formation and maturation. Therefore, it is being reasoned that the characterization of protein – protein interactions involved in microbial biofilms are potential target for development of anti – biofilm drugs.

With regards to the biofilm formation by model organism *Bacillus subtilis*, it has been widely reported and acknowledged that the interaction of ALP protein (TasA) with accessory protein (TapA) is essential for biofilm formation and maturation(Romero et al., 2010; Romero et al., 2011, 2014; Vlamakis et al., 2013); however, information pertaining to the structural characteristics of this protein – protein interaction is quite obscure. The crystal structures of either of these proteins were not available until very recently. The crystal structural of accessory protein TapA is not available till date; therefore an *in vitro* study for characterization of TasA – TapA interaction is fairly difficult. With these rationales, the studies for characterization of TasA - TapA interaction were perused with Computational approaches (e.g. homology modeling; protein-protein interaction prediction with string analysis; protein-protein docking and molecular dynamics simulation etc.). The 3D models of both the proteins were developed and validated. Their stabilities were assessed with MD simulations and the most stable models were subjected to characterization of the interaction sites and identification of the interacting residues amongst the target proteins. Therefore, the protein-protein docking complexes were assessed for TasA_234_-TapA_223_ interaction. A wealth of previous literature shows that hydrogen bonds are vital for providing strength to the proteinprotein interaction complexes(Vakser, 2014); therefore, the hydrogen bonds involved in docked complexes were evaluated. The docking results showed that binding interaction patterns and bond lengths for TasA_234_-TapA_223_ complexes were comparable with the standard values of hydrogen bonds.

The common interacting residues for TasA_234_ – TapA_223_ were identified and characteristically, these interacting amino-acid residues fall within the region of the proteins that are also predicted to be intrinsically disordered. This observation is in agreement with conventional knowledge that suggests for the role of intrinsically disordered regions of different proteins in protein-protein interactions. Another noticeable observation about the identified interfacing residues is that these residues are other than the residues previously reported to have an important role in the TasA-TapA functioning.

The MD simulation of docked protein-protein complexes showed that TasA_234_-TapA_223_ complexes are quite stable. The RMSF values of all the docked complexes showed stability, and only small fluctuations appeared in the graph when assessed during the entire duration of the MD simulation. Similarly, the RMSD values of all the docked complexes also indicated the complexes to be quite stable. On the basis of the results obtained during the present study, it is proposed that we have characterized the critical structural aspect of TasA – TapA interactions and identified the vital amino acid residues involved in these interactions.

## Conclusions

During the present study, the properties of the interaction of *Bacillus subtilis* matrix protein TasA and accessory protein TapA (in their respective processed forms) were characterized using computation biology approaches. The structural and dynamic features were also determined. In conclusion, it is proposed that results obtained during the present study have led to the identification of vital interfacing residues that participate in this TasA -TapA interaction and this interaction utilizes the disorder regions for the interactions. Most noticeably, the interfacing residues in TasA – TapA interaction identified during the present study have never been reported earlier. In future studies, these interfacing residues and their interactions could be a target by small molecule inhibitors and/ or interaction disrupting peptide as a potential strategy for developing Anti-*Bacillus* biofilm agent.

## Acknowledgments

This work was supported by financial support received from SERB – DST, Govt. of India under the Young Scientist Program (Grant No. SERB/YS/LS/2013-294). NV acknowledges the financial support offered in form of the Doctoral Fellowship (F./2015-16/NFO-2015-17-OBC-UTT-28124) by UGC, Govt. of India Department of Science and Technology.

## REFERENCE

2. Abbasi, R., Mousa, R., Dekel, N., Amartely, H., Danieli, T., Lebendiker, M., … Chai, L. (2018). The Bacterial Extracellular Matrix Protein TapA Is a Two-Domain Partially Disordered Protein. Chembiochem. doi: 10.1002/cbic.201800634

3. Amadei, A., Linssen, A. B., & Berendsen, H. J. (1993). Essential dynamics of proteins. Proteins, 17(4), 412–425. doi: 10.1002/prot.340170408

4. Bhutani, I., Loharch, S., Gupta, P., Madathil, R., & Parkesh, R. (2015). Structure, dynamics, and interaction of Mycobacterium tuberculosis (Mtb) DprE1 and DprE2 examined by molecular modeling, simulation, and electrostatic studies. PloS one, 10(3), e0119771

5. Branda, S. S., Chu, F., Kearns, D. B., Losick, R., & Kolter, R. (2006). A major protein component of the Bacillus subtilis biofilm matrix. Mol Microbiol, 59(4), 1229–1238. doi: 10.1111/j.1365-2958.2005.05020.x

6. Branda, S. S., Vik, S., Friedman, L., & Kolter, R. (2005). Biofilms: the matrix revisited. Trends Microbiol, 13(1), 20–26. doi: 10.1016/j.tim.2004.11.006

7. Chapman, M. R., Robinson, L. S., Pinkner, J. S., Roth, R., Heuser, J., Hammar, M., … Hultgren, S. J. (2002). Role of Escherichia coli curli operons in directing amyloid fiber formation. Science, 295(5556), 851–855. doi: 10.1126/science.1067484

8. Claessen, D., Rink, R., de Jong, W., Siebring, J., de Vreugd, P., Boersma, F. G., … Wosten, H. A. (2003). A novel class of secreted hydrophobic proteins is involved in aerial hyphae formation in Streptomyces coelicolor by forming amyloid-like fibrils. Genes Dev, 17(14), 1714–1726. doi: 10.1101/gad.264303

9. Coordinators, N. R. (2013). Database resources of the National Center for Biotechnology Information. Nucleic Acids Res, 41(Database issue), D8–D20. doi: 10.1093/nar/gks1189

10. Costerton, J. W., Lewandowski, Z., Caldwell, D. E., Korber, D. R., & Lappin-Scott, H. M. (1995). Microbial biofilms. Annu Rev Microbiol, 49, 711–745. doi: 10.1146/annurev.mi.49.100195.003431

11. DeLano, W. L. (2002). PyMOL.

12. Diehl, A., Roske, Y., Ball, L., Chowdhury, A., Hiller, M., Molière, N., … Oschkinat, H. (2018). Structural changes of TasA in biofilm formation of <em>Bacillus subtilis</em>. Proceedings of the National Academy of Sciences. doi: 10.1073/pnas.1718102115

13. Dueholm, M. S., Petersen, S. V., Sonderkaer, M., Larsen, P., Christiansen, G., Hein, K. L., … Otzen, D. E. (2010). Functional amyloid in Pseudomonas. Mol Microbiol, 77(4), 1009–1020. doi: 10.1111/j.1365-2958.2010.07269.x

14. Eswar, N., Webb, B., Marti-Renom, M. A., Madhusudhan, M. S., Eramian, D., Shen, M. Y., … Sali, A. (2006). Comparative protein structure modeling using Modeller. Curr Protoc Bioinformatics, Chapter 5, Unit-5 6. doi: 10.1002/0471250953.bi0506s15

15. Flemming, H.-C., & Wingender, J. (2010). The biofilm matrix. Nature Reviews Microbiology, 8, 623. doi: 10.1038/nrmicro2415

16. Flemming, H. C., Neu, T. R., & Wozniak, D. J. (2007). The EPS matrix: the “house of biofilm cells”. J Bacteriol, 189(22), 7945-7947. doi: 10.1128/JB.00858-07

17. Halgren, T. A. (2009). Identifying and Characterizing Binding Sites and Assessing Druggability. Journal of Chemical Information and Modeling, 49(2), 377-389. doi: 10.1021/ci800324m

18. Holton, S. J., Anandhakrishnan, M., Geerlof, A., & Wilmanns, M. (2013). Structural characterization of a D-isomer specific 2-hydroxyacid dehydrogenase from Lactobacillus delbrueckii ssp. bulgaricus. J Struct Biol, 181(2), 179–184. doi: 10.1016/j.jsb.2012.10.009

19. Humphrey, W., Dalke, A., & Schulten, K. (1996). VMD: visual molecular dynamics. J Mol Graph, 14(1), 33–38, 27-38

20. Ichiye, T., & Karplus, M. (1991). Collective motions in proteins: a covariance analysis of atomic fluctuations in molecular dynamics and normal mode simulations. Proteins, 11(3), 205–217. doi: 10.1002/prot.340110305

21. Jamal, M., Ahmad, W., Andleeb, S., Jalil, F., Imran, M., Nawaz, M. A., … Kamil, M. A. (2018). Bacterial biofilm and associated infections. Journal of the Chinese Medical Association, 81(1), 7–11

22. Jefferson, K. K. (2004). What drives bacteria to produce a biofilm? FEMS Microbiol Lett, 236(2), 163–173. doi: 10.1016/j.femsle.2004.06.005

23. Kolter, R., & Greenberg, E. P. (2006). The superficial life of microbes. Nature, 441, 300. doi: 10.1038/441300a

24. Kozakov, D., Hall, D. R., Xia, B., Porter, K. A., Padhorny, D., Yueh, C., … Vajda, S. (2017). The ClusPro web server for protein-protein docking. Nat Protoc, 12(2), 255–278. doi: 10.1038/nprot.2016.169

25. Laskowski, R. A., MacArthur, M. W., Moss, D. S., & Thornton, J. M. (1993). PROCHECK: a program to check the stereochemical quality of protein structures. Journal of Applied Crystallography, 26(2), 283–291. doi: doi:10.1107/S0021889892009944

26. Lemon, K. P., Earl, A. M., Vlamakis, H. C., Aguilar, C., & Kolter, R. (2008). Biofilm development with an emphasis on Bacillus subtilis. Curr Top Microbiol Immunol, 322, 1–16

27. Lynch, A. S., & Abbanat, D. (2010). New antibiotic agents and approaches to treat biofilm-associated infections. Expert Opin Ther Pat, 20(10), 1373–1387. doi: 10.1517/13543776.2010.505923

28. Mashiach, E., Schneidman-Duhovny, D., Andrusier, N., Nussinov, R., & Wolfson, H. J. (2008). FireDock: a web server for fast interaction refinement in molecular docking. Nucleic Acids Res, 36(Web Server issue), W229–232. doi: 10.1093/nar/gkn186

29. Mielich-Süss, B., & Lopez, D. (2015). Molecular mechanisms involved in B acillus subtilis biofilm formation. Environmental microbiology, 17(3), 555–565

30. Negi, S. S., Schein, C. H., Oezguen, N., Power, T. D., & Braun, W. (2007). InterProSurf: a web server for predicting interacting sites on protein surfaces. Bioinformatics, 23(24), 3397–3399

31. Pierce, B. G., Wiehe, K., Hwang, H., Kim, B. H., Vreven, T., & Weng, Z. (2014). ZDOCK server: interactive docking prediction of protein-protein complexes and symmetric multimers. Bioinformatics, 30(12), 1771–1773. doi: 10.1093/bioinformatics/btu097

32. Prism, G. (1994). Graphpad software. San Diego, CA, U S A

33. Pronk, S., Páll, S., Schulz, R., Larsson, P., Bjelkmar, P., Apostolov, R., … Lindahl, E. (2013). GROMACS 4.5: a high-throughput and highly parallel open source molecular simulation toolkit. Bioinformatics, 29(7), 845–854. doi: 10.1093/bioinformatics/btt055

34. Romero, D., Aguilar, C., Losick, R., & Kolter, R. (2010). Amyloid fibers provide structural integrity to Bacillus subtilis biofilms. Proc Natl Acad Sci USA, 107(5), 2230–2234. doi: 10.1073/pnas.0910560107

35. Romero, D., Vlamakis, H., Losick, R., & Kolter, R. (2011). An accessory protein required for anchoring and assembly of amyloid fibres in B. subtilis biofilms. Mol Microbiol, 80(5), 1155–1168. doi: 10.1111/j.1365-2958.2011.07653.x

36. Romero, D., Vlamakis, H., Losick, R., & Kolter, R. (2014). Functional analysis of the accessory protein TapA in Bacillus subtilis amyloid fiber assembly. J Bacteriol, 196(8), 1505–1513. doi: 10.1128/JB.01363-13

37. Serrano, M., Zilhao, R., Ricca, E., Ozin, A. J., Moran, C. P., Jr., & Henriques, A. O. (1999). A Bacillus subtilis secreted protein with a role in endospore coat assembly and function. J Bacteriol, 181(12), 3632–3643

38. Sievers, F., Wilm, A., Dineen, D., Gibson, T. J., Karplus, K., Li, W., … Higgins, D. G. (2011). Fast, scalable generation of high-quality protein multiple sequence alignments using Clustal Omega. Mol Syst Biol, 7, 539. doi: 10.1038/msb.2011.75

39. Szklarczyk, D., Morris, J. H., Cook, H., Kuhn, M., Wyder, S., Simonovic, M., … von Mering, C. (2017). The STRING database in 2017: quality-controlled protein-protein association networks, made broadly accessible. Nucleic Acids Res, 45(D1), D362–D368. doi: 10.1093/nar/gkw937

40. Taglialegna, A., Lasa, I., & Valle, J. (2016). Amyloid Structures as Biofilm Matrix Scaffolds. J Bacteriol, 198(19), 2579–2588. doi: 10.1128/JB.00122-16

41. Terra, R., Stanley-Wall, N. R., Cao, G., & Lazazzera, B. A. (2012). Identification of Bacillus subtilis SipW as a bifunctional signal peptidase that controls surface-adhered biofilm formation. J Bacteriol, 194(11), 2781–2790. doi: 10.1128/JB.06780-11

42. Vakser, I. A. (2014). Protein-protein docking: from interaction to interactome. Biophys J, 107(8), 1785–1793. doi: 10.1016/j.bpj.2014.08.033

43. Vangone, A., Spinelli, R., Scarano, V., Cavallo, L., & Oliva, R. (2011). COCOMAPS: a web application to analyze and visualize contacts at the interface of biomolecular complexes. Bioinformatics, 27(20), 2915–2916. doi: 10.1093/bioinformatics/btr484

44. Vlamakis, H., Chai, Y., Beauregard, P., Losick, R., & Kolter, R. (2013). Sticking together: building a biofilm the Bacillus subtilis way. Nature Reviews Microbiology, 11(3), 157

45. Xue, B., Dunbrack, R. L., Williams, R. W., Dunker, A. K., & Uversky, V. N. (2010). PONDR-FIT: a meta-predictor of intrinsically disordered amino acids. Biochim Biophys Acta, 1804(4), 996–1010. doi: 10.1016/j.bbapap.2010.01.011

