## Supplementary figures and images for "Computational investigation for modelling the protein-protein interaction of TasA_(28-261)_ – TapA_(33-253)_: a decisive process in biofilm formation by *Bacillus subtilis*"

### Supp Fig 1

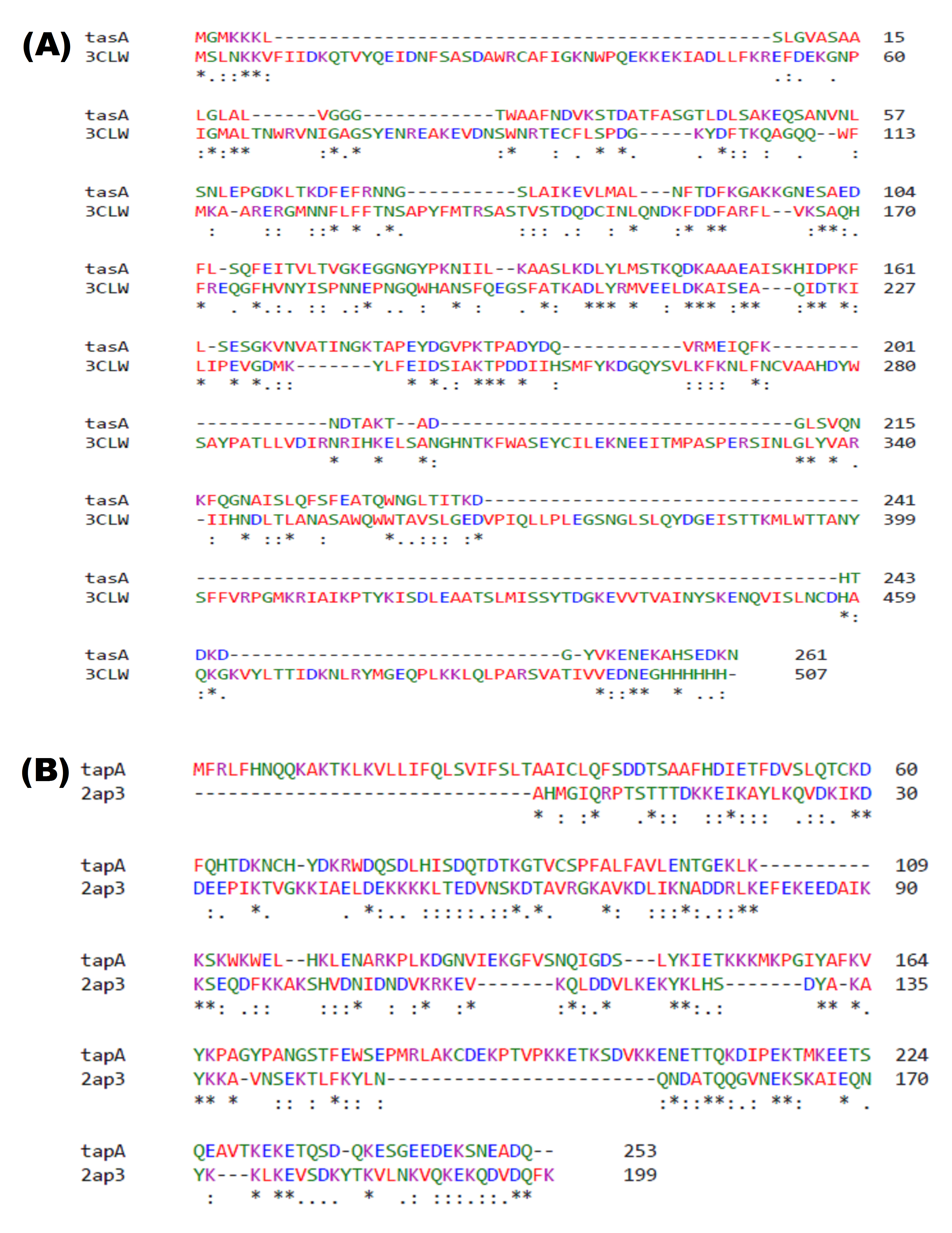

### Supp Fig 2

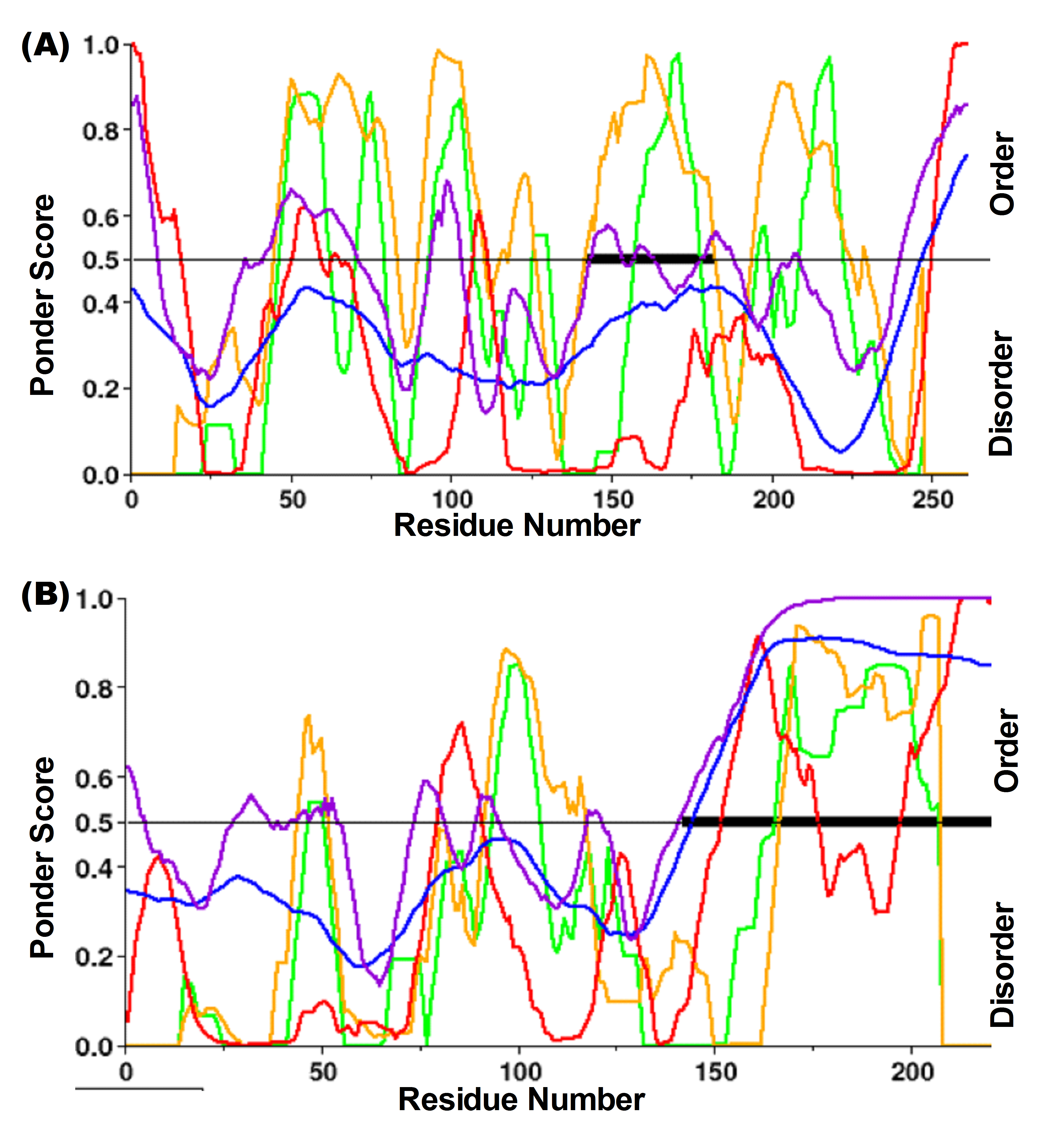

### Supp Fig 3

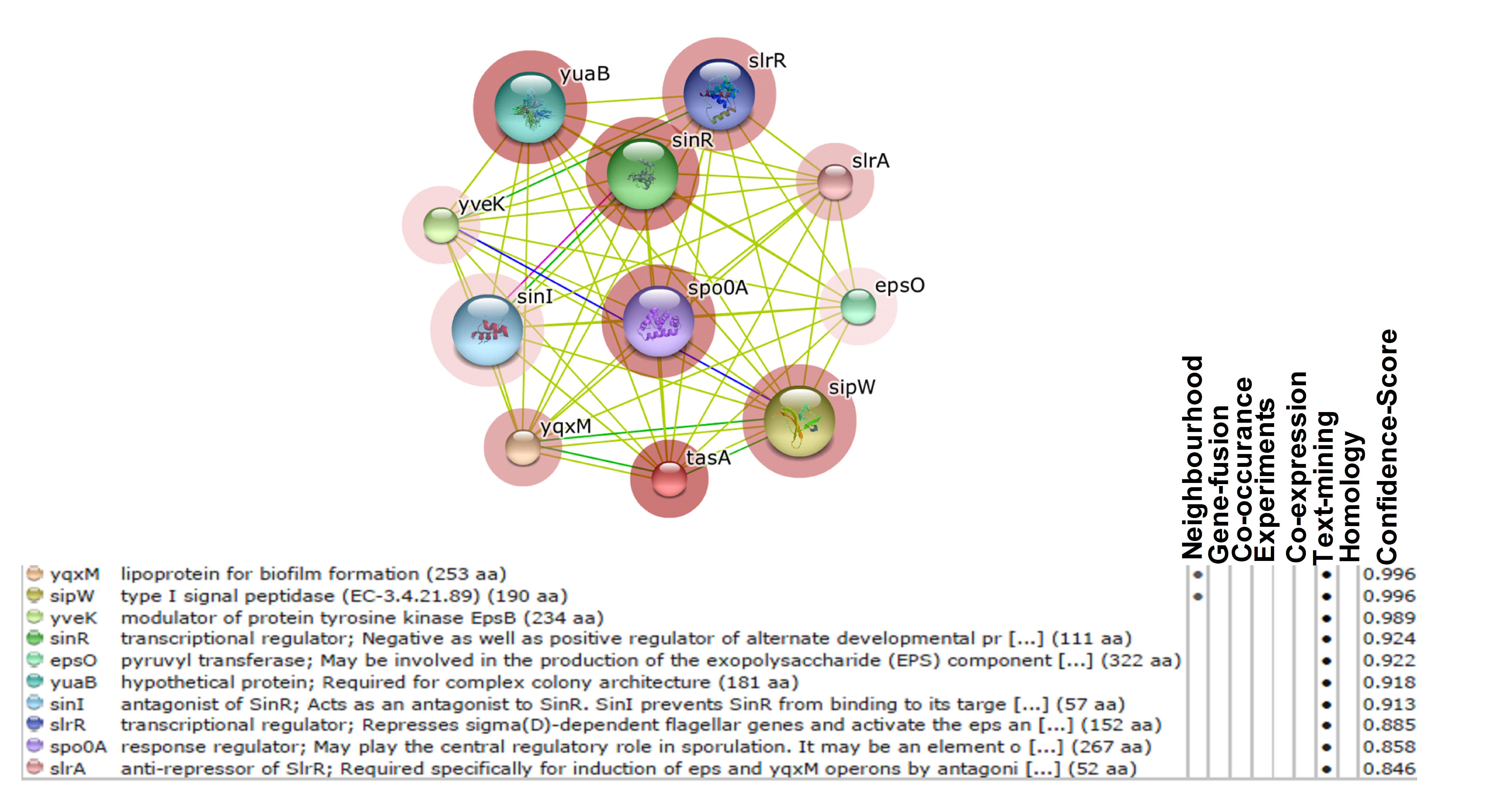
